# The genome-wide rate and spectrum of spontaneous mutations differs between haploid and diploid yeast

**DOI:** 10.1101/250365

**Authors:** Nathaniel P. Sharp, Linnea Sandell, Christopher G. James, Sarah P. Otto

## Abstract

By altering the dynamics of DNA replication and repair, alternative ploidy states may experience different rates and types of new mutations, leading to divergent evolutionary outcomes. We report the first direct comparison of the genome-wide spectrum of spontaneous mutations arising in haploid and diploid forms of the budding yeast. Characterizing the number, types, locations, and effects of thousands of mutations revealed that haploids were more prone to single-nucleotide and mitochondrial mutations, while larger structural changes were more common in diploids. Mutations were more likely to be detrimental in diploids, even after accounting for the large impact of structural changes, contrary to the prediction that diploidy masks the effects of recessive alleles. Haploidy is expected to reduce the opportunity for conservative DNA repair involving homologous chromosomes, increasing the insertion-deletion rate, but we found little support for this idea. Instead, haploids were more susceptible to particular single-nucleotide mutations in late-replicating genomic regions, resulting in a ploidy difference in the spectrum of substitutions. In diploids we detect mutation rate variation among chromosomes in association with centromere location, a finding that is supported by published polymorphism data. Diploids are *Saccharomyces cerevisiae* not simply doubled haploids; instead, our results predict that the spectrum of spontaneous mutations will substantially shape the dynamics of genome evolution in haploid and diploid populations.

## Introduction

Mutations play a critical role in evolution and adaptation, but these spontaneous genetic changes are often hazardous, reducing the health of individuals and imposing a genetic load on populations. While individual mutations are chance events, the biological processes that produce or prevent mutation have the potential to vary among genetic contexts, leading to consistent biases in the numbers, locations, and types of genetic changes that occur. As a consequence, populations with alternative genome architectures may have access to different kinds of genetic variation and may be subject to different risks of deleterious events. Characterizing variation in the mutation process is therefore important to understanding why and how genome architecture evolves, interpreting neutral and adaptive sequence evolution, and predicting future genetic change.

One aspect of genome architecture that has broad potential to affect the mutation process is ploidy state, the number of homologous chromosome sets per cell. Ploidy varies among cells as a consequence of meiosis and syngamy and can also vary among individuals, e.g., in haplodiploid organisms. Transitions in predominant ploidy level have occurred many times across the tree of life (1, 2), but the evolutionary and ecological drivers and consequences of these changes are still unclear.

All else being equal, the genome-wide mutation rate of diploid cells should be twice that of haploid cells, as they have twice as many nucleotide sites with the potential to mutate. However, ploidy level is likely to have additional effects on the rate and spectrum of mutations. The presence of a homologous chromosome template allows DNA double-strand breaks (DSBs) to be repaired using conservative homology-directed pathways rather than competing error-prone end-joining pathways that generate insertion-deletion (indel) mutations (3). The continual access to homologous template DNA in diploid cells may therefore reduce the rate of indels relative to haploids. While diploidy may reduce susceptibility to certain forms of DNA damage, the presence of multiple genome copies may also increase the likelihood of spontaneous structural changes and rearrangements through non-homologous crossing over (3). There are therefore reasons to expect differences in the mutational properties of alternative ploidy states.

Estimates of genome-wide mutation rates are available from mutation accumulation (MA) experiments with the budding yeast *S. cerevisiae*, which can grow mitotically in either a haploid or diploid state. In these experiments, replicate lines are periodically subjected to single-cell bottlenecks, allowing mutations with mild and moderate effects to accumulate as though they were selectively neutral. Previous studies have examined either haploids or diploids, preventing direct comparison.

Genome-wide patterns of spontaneous mutation have been somewhat better characterized in diploid *S. cerevisiae* (4, 5) than in haploids, where sampling has been limited (6, 7). At face value, previous estimates suggest that the rate of single-nucleotide mutations (SNMs) per base pair may be greater in haploids than in diploids, and it has been proposed that yeast genomes are more stable in the diploid state (4, 5). However, these different inferred rates could reflect variation among lab strains, methodology, or environments. Additionally, in a previous study of haploids, all four lines examined by genome sequencing apparently became diploid spontaneously during the experiment (6), complicating inference of the mutation rate in haploid cells.

To formally investigate the impact of ploidy on the mutation process, we accumulated mutations in isogenic replicate haploid and diploid lines of *S. cerevisiae* derived from a common haploid ancestor. We maintained 220 lines under relaxed selection by enforcing single-cell bottlenecks every day (∼16 generations) for 100 days, for a total of ∼336,000 generations across the experiment. To address the possible role of homology-directed DSB repair, we deleted the gene *RDH54* / *Tid1* in the ancestor of half of the lines. There is evidence that this gene is essential for mitotic recombinational repair between homologous chromosomes in diploids, but not between sister chromatids in haploids or diploids (8). Genome sequencing of these lines revealed over 2000 mutation events, with clear impacts of ploidy on the rate, spectrum, locations, and fitness consequences of spontaneous mutations, but little evidence that homology-directed DSB repair is a major factor in these differences.

## Results

### Growth Rates and Ploidy

To account for potential differences among treatments in rates of cell division, we measured the number of cells per colony in the MA lines throughout the bottlenecking procedure and used these values to calculate mutation rates per cell division (Table 1). We found small but significant treatment effects on growth rate, with a more rapid decline in diploid lines than in haploid lines over the course of the experiment (Fig. 1A), 1B).

**Table 1.** Sample sizes.

|  | Haploid |  | Diploid |  |
| --- | --- | --- | --- | --- |
|  | *RDH54+* | *rdh54Δ* | *RDH54+* | *rdh54Δ* |
| MA lines | 53 | 53 | 63 | 51 |
| Cell divisions/transfer | 15.7 | 15.2 | 15.8 | 15.7 |
| Transfers/line* | 94.9 | 97.5 | 100.0 | 98.9 |
| Median coverage/site/line (range†) | 23 (16, 30) | 22 (10, 35) | 44 (20, 65) | 44 (30, 62) |
| Callable Nuc sites/line‡ | 11296337 | 11255081 | 23088492 | 22684822 |
| Callable Mt sites/line | 72036 | 70426 | 74868 | 75028 |

Nuc, nuclear; Mt, mitochondrial.
\*Some lines were re-initialized partway through the procedure and therefore underwent <100 transfers as unique lines.
†Min. and max. of median coverage among lines.
‡Incorporates ploidy, including ancestral trisomy of chromosome XI in some lines.

**Figure 1.**
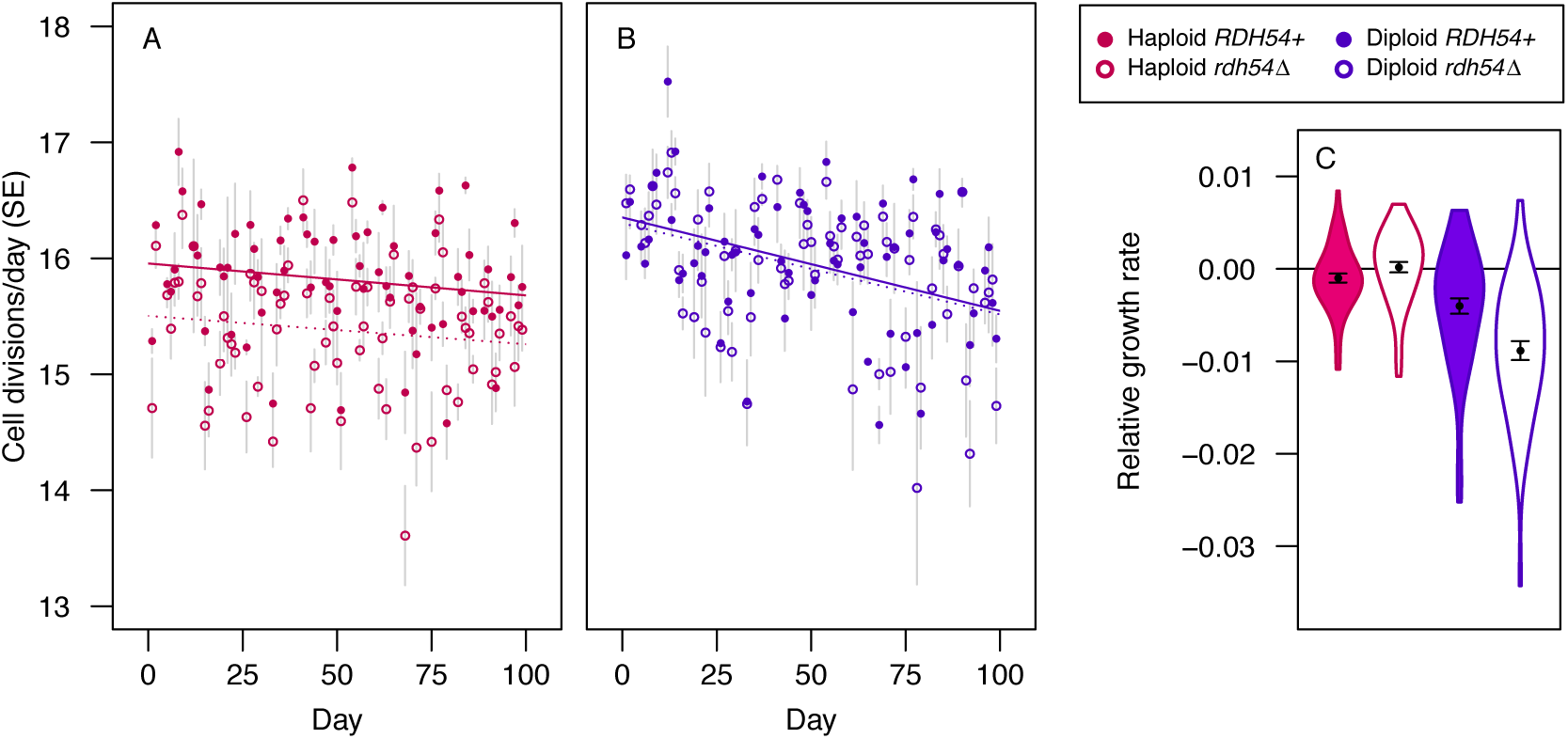
Cell divisions during MA and growth rates following MA. Cell divisions per day (haploids A, diploids B) depended on the interaction of ploidy level, *RDH54* status, and time (>3200 colonies measured; linear model: ploidy × *RDH54*: *P* < 10^−8^; ploidy × time: *P* < 0.01). Solid (dashed) lines show linear regression for *RDH54* + (*rdh54*Δ) treatments. (C) Violin plots of maximum growth rates after 100 transfers relative to ancestral controls, accounting for block effects. Points (error bars) show means (SEs). Steeper declines in cell divisions per day in diploids relative to haploids during MA (A and B) are consistent with relative growth rate estimates following MA (linear mixed model based on 4000 growth curves: mutation accumulation × ploidy × *RDH54* : *P* < 0.05). Diploid MA lines have significantly reduced growth rates relative to the ancestor (MA: *P* < 10^−6^, MA × *RDH54* : *P* < 0.05), unlike haploid lines (MA: *P* = 0.64). See Dataset S2 for data on growth rates following MA.

After 100 bottlenecks we measured the growth rates of MA lines and ancestral controls in liquid culture, revealing that diploid MA lines, but not haploid MA lines, show reduced growth relative to ancestral controls (Fig. 1C), consistent with the measures of colony growth on plates during MA. All four groups of MA lines (ploidy × RDH54 status) show significant genetic variance for growth rate (all χ^2^ > 24.8, all *P* < 10^−6^), whereas there was no significant genetic variance detected in any group of ancestral control lines (all χ^2^ < 2.6, all *P* > 0.11). The finding that diploid but not haploid lines show reduced growth following MA is surprising, as the effects of recessive deleterious mutations would be masked in diploids. Below we discuss this result in relation to the genomic data (see *Growth rates in relation to mutations*).

Haploid yeast are known to become diploid spontaneously (e.g., through defects in mitosis), and diploids can similarly change ploidy level. Such changes are commonly observed in yeast evolution experiments with large populations, but the spontaneous rate of ploidy change per cell division is unclear. Using flow cytometry, we found that all MA lines retained their original ploidy level at the end of the experiment (Dataset S1; confidence interval, CI, for rate of change from haploid to diploid: 0 to 2.3 × 10^−5^ per cell division; see below for results on aneuploidy). This is in contrast to a previous finding (6) where four out of four haploid MA lines became diploid over the course of ∼4800 generations (rate [95% CI]: 20.8 [5.7, 53.3]× 10^−5^). This difference suggests either that the higher frequency of single-cell bottlenecks in our experiment (daily rather than every 3−4 days) reduced the opportunity for selection favouring diploidy or that yeast strains vary in their propensity to spontaneously change ploidy.

We observed changes in the mating behaviour of three MA lines, and describe the likely genetic basis for these changes in *SI Text*.

### Point Mutations

We detected >2000 point mutations (Dataset S2). Mutation rates, accounting for numbers of cell divisions and callable sites, are shown in Fig. 2). There was no effect of mating type on mutation rates within haploids (Fig. S1), and so we pooled the data for haploid MATá and MAT α lines throughout our analyses. We detected a significant effect of ploidy level on the rate of single-nucleotide mutations (SNMs) (binomial test: *P* < 10^−11^), with no effect of *RDH54* status (*P* > 0.24 in either ploidy level). The per-nucleotide SNM rate was 40% higher in haploids (4.04 × 10^−10^, 95% CI: 3.75 to 4.34 × 10^−10^) than in diploids (2.89 × 10^−10^, 95% CI: 2.73 to 3.06 × 10^−10^). Thus, although diploids have twice as many nucleotide sites as haploids they incur only 1.43 times as many SNMs per cell division.

**Figure 2.**
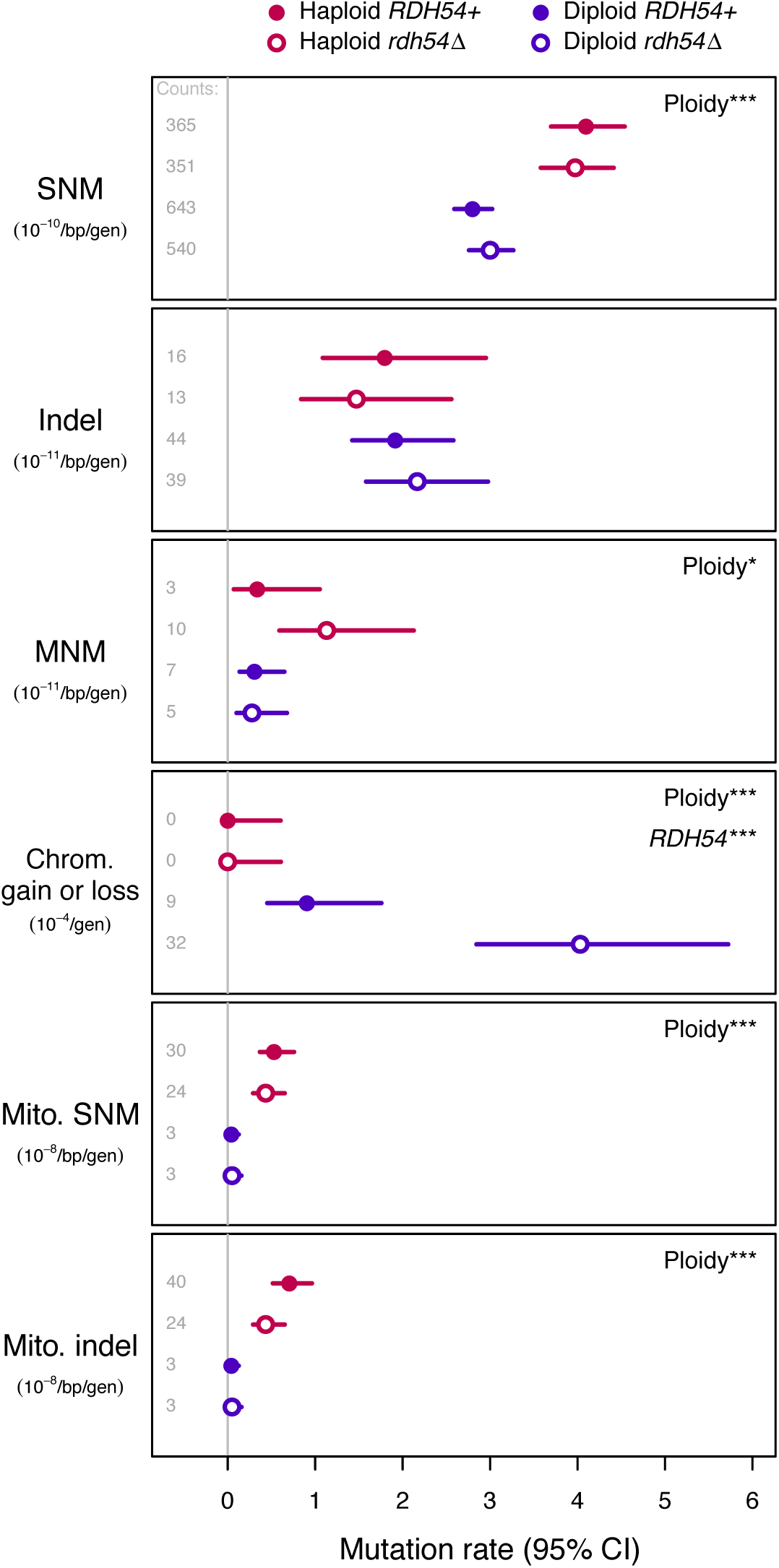
Mutation rates in each group of MA lines, with 95% confidence intervals. Panels show single nucleotide mutations (SNM), insertions and deletions (indel), multinucleotide mutations (MNM), chromosome gains and losses, mitochondrial SNM, and mitochondrial indels. Rates represent events per base pair per generation except for whole-chromosome gains and losses (events per generation). For mitochondrial events we consider all treatments effectively haploid. The absolute numbers of events observed (“counts”) are given at the left of each panel; note that detection power differs among groups such that rates are not necessarily proportional to mutation counts. Statistically significant treatment effects are noted at the right of each panel (binomial tests accounting for detection power; “*” *P* < 0.05, “**” *P* 0.01, “***” *P* < 0.001). See Dataset S2 for a numeric summary of rates and confidence intervals.

Because diploidy provides greater access to homologous DSB repair we expected to see a higher rate of indel mutations in haploids and in diploids lacking the *RDH54* gene. However, the indel rate did not depend on ploidy level (binomial test: *P* = 0.52) or *RDH54* status (*P* > 0.58 in either ploidy level; Fig. 2). In fact, our point estimate of the per-nucleotide rate of indels is 24% greater in diploids (2.03 × 10^−11^) than in haploids (1.63 × 10^−11^), such that the indel rate per cell division is 2.48 times higher in diploids. There was a non-significant tendency towards deletions among indel events (62 deletions and 50 insertions; binomial test: *P* = 0.30).

We considered whether selection during the experiment might have affected rates of mutation accumulation. Accounting for power to detect mutations in genic and non-genic regions and assuming that genic and non-genic regions are equally likely to mutate, there is no evidence that SNMs were less likely to occur in genes than expected (observed 74.0%, expected 73.9%, binomial test: *P* = 0.92) or less likely to be non-synonymous than expected (observed 76.7%, expected 76.1%; binomial test: *P* = 0.62), with no difference in these rates between ploidy or *RDH54* levels. There is therefore no evidence that the accumulation of SNMs was influenced by selection in our experiment.

By contrast, indels were found in genes less often than expected (observed 46.4%, expected 73.9%; binomial test: *P* < 10^−9^), as observed in other datasets of this kind (5). Selection acting more effectively against indels in haploids could mask a true difference in indel rate. However, the fraction of indels that accumulated in genes does not differ significantly between ploidy levels (Fisher’s exact test: odds ratio 1.10, *P* = 0.83), and there is no indication that non-genic indel rates differed by RDH54 status or ploidy (Fig. S2). It is therefore not apparent that the pattern of indel accumulation among treatment groups in our experiment was driven by selection.

As in previous studies where MA lines have been sequenced, we find evidence for some substitutions occurring in close proximity to one another, which we categorize as multi-nucleotide mutations (MNMs; (9). As with SNMs, the MNM rate was significantly higher in haploids (0.73 × 10^−11^;) than diploids (0.29 × 10^−11^; Fig. 2); binomial test: *P* < 0.05), suggesting that replication errors associated with ploidy generate both SNMs and MNMs.

We identified 38 homozygous SNMs and one homozygous indel in diploid lines, reflecting loss of heterozygosity (LOH) events that can occur through mitotic crossing over, non-crossover gene conversion, or chromosome loss and duplication (3, 10). The rate of homozygous mutations did not differ between *RDH54* treatments (binomial test: = 0.63). For six homozygous mutations another mutation occurred distally on the same chromosome arm in the same MA line, and of these, three were also homozygous, a higher rate than expected if all LOH events occurred independently (binomial test, *P* < 0.001). These pairs of homozygous mutations were separated by 132−451 kb, suggesting that a substantial fraction of LOH events are due to mitotic crossing over or chromosome loss/duplication rather than relatively short gene conversion tracts.

### Large-Scale Mutations

We used sequencing coverage across the genome to detect changes in chromosome copy number (aneuploidy), which result from mitotic nondisjunction. 49 out of 63 *RDH54* + diploid lines were found to carry three copies of chromosome XI, which we established to be an ancestral polymorphism in that group of lines (*Materials and Methods*). We also observed 41 *de novo* chromosome gains and losses, all within the diploid MA lines (Fig. 2). Chromosome gains (35 cases of trisomy and 2 cases of tetrasomy) were found more often than chromosome losses (4 losses, including a loss of a trisomic chromosome XI restoring euploidy; binomial test: *P* < 10^−6^; Fig. 3A), as observed in a previous MA experiment (5). While the rarity of chromosome losses relative to gains may reflect stronger selection against monosomy in diploids or a higher chance of reversion to euploidy from monosomy in MA experiments, our observed rates of chromosome loss are generally consistent with previous estimates obtained using other methods (reviewed in (3). The derived versus ancestral allele depths for substitutions conformed well to the expected patterns on aneuploid and non-aneuploid chromosomes (Fig. S3), confirming that the chromosome XI trisomy occurred prior to MA and that other aneuploidies occurred uniformly throughout MA.

**Figure 3.**
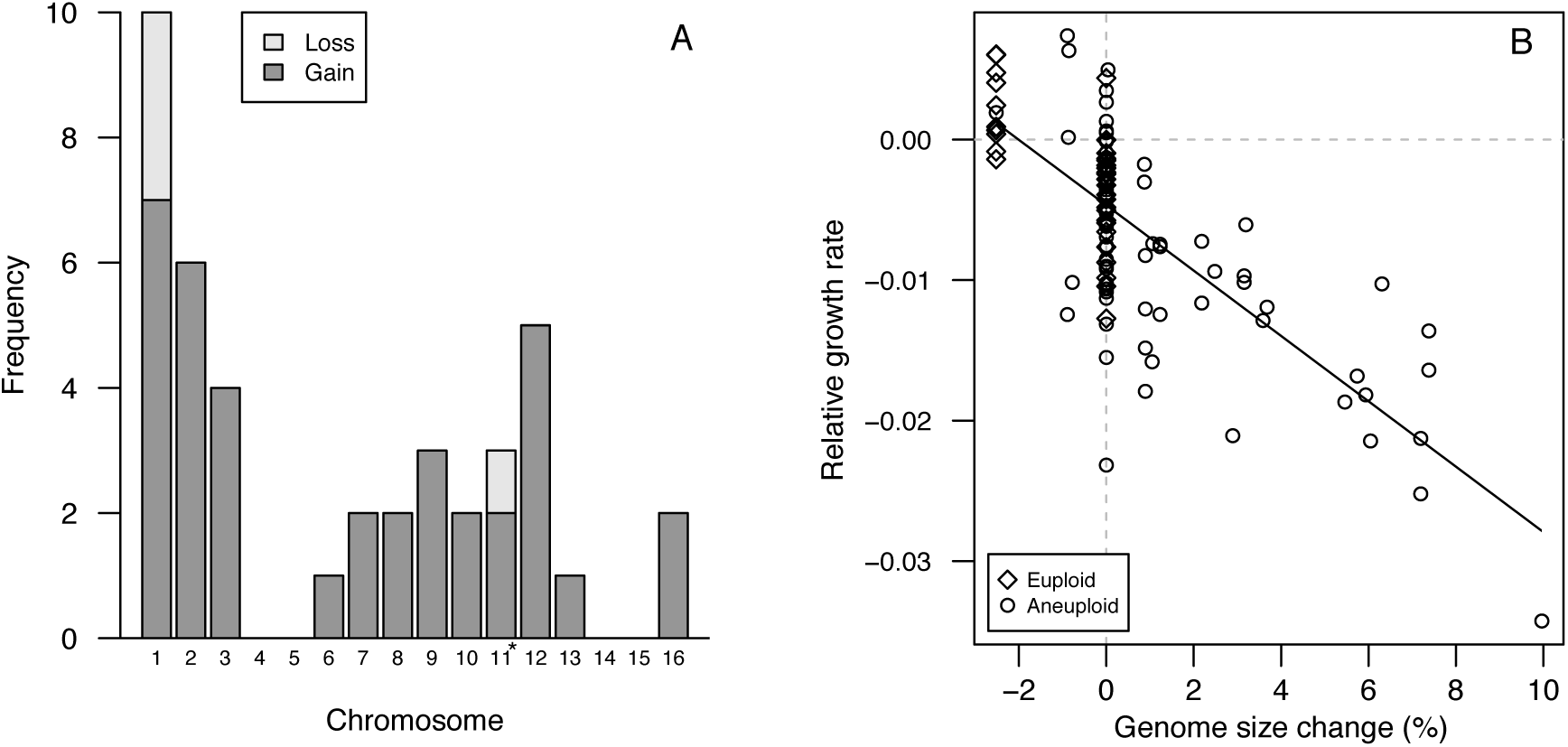
Gains and losses of each chromosome during MA and the fitness consequences of aneuploidy. (A) Gains were more common than losses (binomial test, *P* < 10^−6^), and rates vary among chromosomes (χ^2^ = 42.1, *P* < 0.001). Several + diploids were determined to be ancestral and are not scored as gains; in one case the extra copy was lost, restoring euploidy, and this is scored here as a loss. See Dataset S2 for aneuploid line IDs. (B) Genome size relative to controls was negatively correlated with maximum growth rate relative to controls in diploid MA lines (*r* = −0.74, *df* = 112, *P* < 10^−15^). Diploid cases of trisomy for chromosome 11 in *RDH54* + controls had trisomy for chromosome 11, so MA lines with only this trisomy are scored as having zero genome size change and MA lines without this trisomy were scored as having a genome size reduction unless other aneuploidy was present. This correlation persists when only aneuploid lines are considered (*r* = − 0.70, *df* = 78, *P* < 10^−12^) or when lines with no change are excluded (*r* = −0.84, *df* = 45, *P* < 10^−12^).

While the rate of aneuploidy in *RDH54* + diploids (0.90 × 10^−4^) is very similar to previous observations from diploid yeast ((5); rate 1.04 × 10^−4^), we found that *rdh54* Δ elevated the rate of aneuploidy 4-fold overall and 4.5-fold within diploids (Fig. 2); binomial test: *P* < 10^−4^), indicating that *RDH54* plays a role in mitotic chromosome segregation in addition to its role in meiosis (8, 11), as suggested previously (12). Given the absence of aneuploidy in haploids, we cannot determine whether the increase in aneuploidy in *rdh54* Δ lines also applies to haploids or is unique to diploids.

We find that the rate of aneuploidy is significantly higher in diploids on a per-cell-division basis (Fig. 2); binomial test: *P* < 10^−11^). This pattern persists when excluding chromosome losses, which would be lethal in haploids (binomial test: *P* < 10^−10^), and when excluding *rdh54* Δ lines (binomial test:*P* < 0.05). Accounting for the number of chromosomes per cell that could potentially increase in copy number (two-fold higher in diploids), the effect of ploidy on the rate of chromosome gains remains significant overall (binomial test: < 10^−5^). Considering only gains in the *RDH54* + lines, however, we cannot reject the hypothesis that the opportunity for nondisjunction is simply two-fold higher in diploids than in haploids (binomial test: = 0.20).

The probability of aneuploidy was not uniform across chromosomes (Fig. 3A), and there was a negative but non-significant correlation between the rate of gain or loss and chromosome length (*r*= −0.41, *P* = 0.11), supporting the view that chromosome size is not the sole determinant of aneuploidy rates (3, 5). Variation in aneuploidy rates among chromosomes in our experiment was correlated with aneuploidy rates observed previously in diploids exhibiting chromosome instability (13) (*r*= 0.57 *P*<0.05). As in this previous study (13), we found that growth rates were highly correlated with genome size among aneuploid lines (Fig. 3B).

In addition to aneuploidy, coverage profiles revealed three cases of large segmental duplications, all heterozygous in diploids (Fig. S4), ranging in size from 17 kb to 211 kb. The approximate breakpoints of all three duplications are closer to known Ty elements (mean distance 700 bp, range 100−1662 bp) than expected by chance (simulation of 10^4^ sets of 6 breakpoints; mean distance to Ty ≤ 700: *P* < 10^−^), supporting the view that Ty elements often anchor large duplications in yeast (14–16). Our estimate of the large duplication rate in diploids is 1.68 [95% CI: 0.34, 5.24] × 10^−5^ per cell division, consistent with a previous observation of diploid MA lines ((5) approximately 1.00 × 10^−5^).

The presence of mutations could increase the subsequent mutation rate by directly altering DNA repair genes, by increasing genome instability (3, 7), or if genetic quality affects DNA repair (17). However, we found no evidence for an effect of aneuploidy on the point mutation rate and no evidence for overdispersion in the numbers of mutations per line (*SI Text*
).

### Mitochondrial Mutations

The cellular environment, replication mechanisms and nucleotide composition of mitochondrial (mt) genomes are distinct from those of nuclear genomes, but it is not clear how nuclear genome ploidy might interact with mt mutation. We found that the rate of mt mutation was 11.8 times higher in haploids (SNM: 4.82 × 10^−9^, indel: 5.71 × 10^−9^) than in diploids (SNM: 4.47 × 10^−10^, 10^−10^; Fig. 2). Mt mutation rate estimates based on fewer events from a previous study of four haploid MA lines (which became diploid during MA) are higher than the haploid rates we observed ((6); SNM: 12.2 × 10^−9^, indel: 10.4 × 10^−9^). indel: 4.47 × As in this previous study we found that the haploid substitution rate was higher in mt than in the nuclear genome (11.9-fold difference, binomial test: *P* < 10^−15^), but this was not the case for diploids (1.6-fold difference, binomial test: *P* = 0.30).

As with nuclear point mutations, there was no evidence that mt SNMs occurred in genes less often than expected (observed: 0.27, expected: 0.35; binomial test, *P* = 0.23), but mt indels were less likely to accumulate in genes than expected (observed: 0.19; binomial test, *P* < 0.01). The fraction of events in genes did not differ between ploidy levels for either SNMs (*G* = 0.32, *P* = 0.57) or indels (*G* = 0.07, *P* = 0.79), and so there is no indication that the elevated rate of mt mutations in haploids relative to diploids is the result of more effective selection against such events in diploids. Small insertions and deletions were about equally prevalent among mt events (37 deletions, 33 insertions; binomial test: *P* = 0.72).

Mt deletions or other mutations can sometimes result in a “petite” phenotype due to defects in respiratory function. Although we attempted to prevent the accumulation of such variants, two haploid lines had a petite phenotype by the end of the experiment. An examination of sequencing coverage in these lines indicated large deletions in COX1 with very similar breakpoints (deleted locations approximately 16438−24230 and 16435−24592). Mt deletions with these approximate breakpoints are known to occur frequently, possibly due to auto-catalytic activity (18). Additionally, two haploid lines showed large mt deletions in non-genic regions based on coverage (deleted locations approximately 4824−5751 and 80559–83041). An additional haploid line with a missense mutation in COX1 also showed poor respiration. Thus, in the mt genome, both point mutations and structural changes were observed more often in haploids, unlike in the nuclear genome where structural changes were observed more often in diploids.

### Spectrum of Nucleotide Changes

Haploids and diploids differed in the spectrum of SNMs (Fig. 4A). In particular, A-to-T and C-to-G changes were more common among haploid SNMs than among diploid SNMs, whereas the reverse was true for C-to-A changes. Overall, mutations at A/T sites made up a larger fraction of SNMs in haploids than in diploids (Fig. 4B), although all six types of substitution occurred more frequently in haploids than in diploids (Fig. S5).

**Figure 4.**
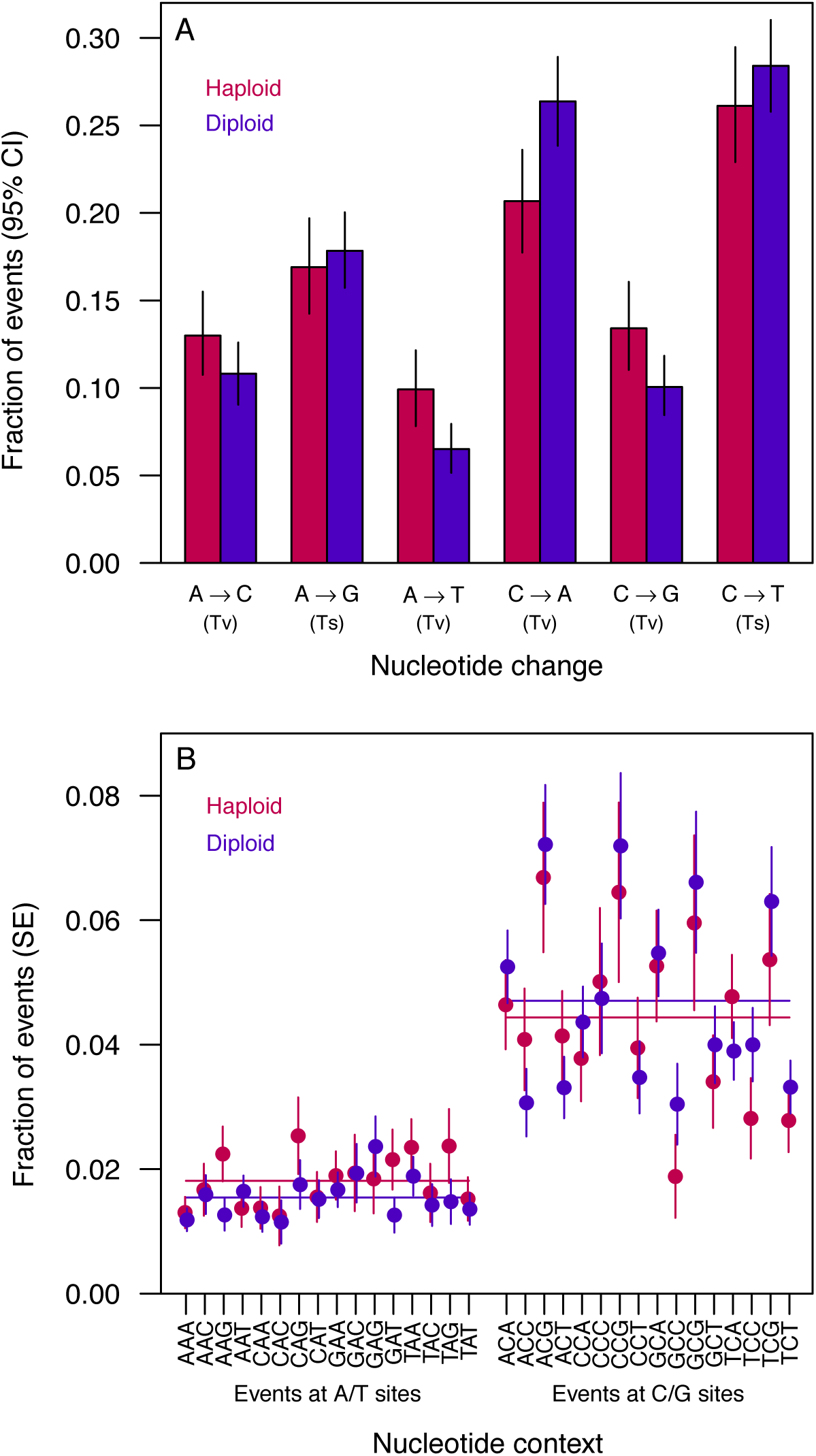
Spectrum of nuclear SNMs. (A) The fraction of SNMs of each type, including complementary changes. Tv = transversion, Ts = transition. status did not significantly affect the SNM spectrum (*G* = 4.5, *P* = 0.48), but the SNM spectrum differed between ploidy levels (*G* = 19.8, < 0.01), particularly among transversion mutations (*G* = 17.9, *P* < 0.001). By contrast, the spectrum of transition mutations and the overall transition-to-transversion ratio did not vary significantly by ploidy (*G* = 0.04, *P* = 0.84; odds ratio = 0.88, *P* = 0.18, respectively). (B) The fraction of SNMs in each 3-bp context, including the complementary context, centered on the focal site, accounting for the frequency of each context in the genome. While the context of SNMs did not differ significantly between haploids and diploids when considering all contexts (*G* = 21.5, *P* = 0.90 Events at A/T sites Events at C/G sites horizontal lines show means across A/T or C/G sites for each ploidy level). The overall rate of SNMs depended on 3-bp context at G/C sites; the fraction of mutations at A/T vs. C/G sites differed between ploidy levels (odds ratio = 1.22, *P* < 0.05; *G* = 81.9, *P* < 10^−10^) but not at A/T sites (*P* = 20.9 *G* = 0.14). The mutation rate in XCG contexts (where X is any base) was elevated relative to the rate for C/G sites in other contexts (binomial test: *P* < 10^−11^).

The transition:transversion ratio did not differ significantly between ploidy levels (haploid: 0.75; diploid 0.86; odds ratio = 0.88, *P* = 0.18), and the overall ratio, 0.82 (95%CI: 0.75 to 0.90) fell between previous estimates for initially-haploid MA lines (0.62, (6) and diploid MA lines (0.95, (5).

SNM rates were affected by adjacent nucleotide context at C/G sites but not A/T sites, with no differences in context effects between ploidy levels (Fig. 4B). In our experiment, both haploids and diploids show an elevated substitution rate at CpG sites compared to C/G sites in other contexts (binomial test, *P* < 10^−11^; Fig. 4B), with a ∼2-fold elevation in the rate of C-to-T mutations in this context (binomial test, *P* < 10^−14^). Although *S. cerevisiae* shows little evidence for cytosine methylation (but see (19), which can increase the frequency of C-to-T mutations in CpG contexts (20), such an increase has been observed in both diploid *S. cerevisiae* (5) and haploid *S. pombe*(21, 22).

### Genomic Locations of Mutations

Previous studies have found evidence for increased mutation in late-replicating genomic regions in haploids (23) and non-significant trends in diploids (5). Here, we use published information on replication across the yeast genome (24) to compare the effect of replication timing between ploidy levels. Replication timing across the genome is very similar between ploidy levels (24), but we find that SNMs were more likely to occur in later-replicating regions in haploids, whereas there was no effect of replication timing on the rate of SNMs in diploids (Fig. 5A). This effect of replication timing in haploids at least partly explains the overall difference in SNM rates between ploidy levels: we found that haploid and diploid SNM rates were similar within early-replicating genomic regions, but diverged in later-replicating regions (Fig. S6).

**Figure 5.**
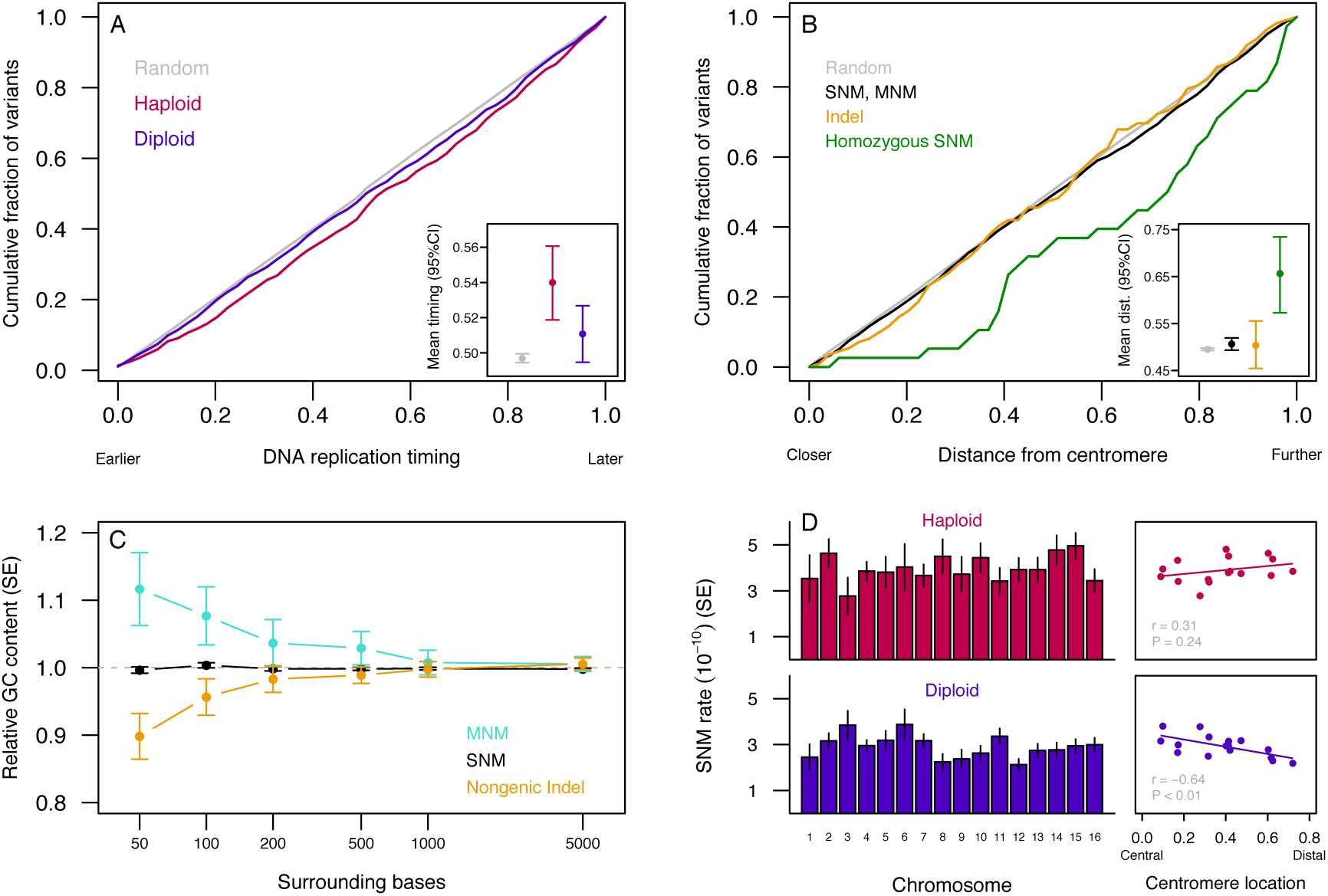
Genomic context of nuclear mutations. (A) The DNA replication timing of mutated sites differed significantly between haploids (red) and diploids (blue) (Wilcoxon test, *P* < 0.05). Haploid mutations arose in later-replicating positions relative to random callable sites (Wilcoxon test, *P* < 10^−4^), whereas diploid mutations did not differ significantly from the random expectation (Wilcoxon test, *P* = 0.09). Inset: mean replication timing for each group. See also Fig. S6. (B) Mutations vs. distance from the centromere, as a fraction of maximum possible distance. All event types except indels occurred further from the centromere than random callable sites (Wilcoxon tests; SNM and MNM: *P* < 0.05, Homozygous: *P* < 0.001), and homozygous diploid SNMs are further from the centromere than heterozygous SNMs (Wilcoxon test: *P* < 0.001). Inset: mean distance from centromere for each event type. (C) GC content surrounding mutations relative to GC content surrounding equivalent random sites (A/T or C/G sites for substitutions; genic or non-genic sites for all mutation types). We consider only non-genic indels here because of possible selection against genic indels. MNM values represent the mean for the component sites. MNMs were associated with higher GC content within 50 bp (bootstrap *P* < 0.05), whereas nongenic indels were associated with lower GC content within 50 bp (bootstrap *P* < 0.01). (D) Variation in SNM rate among chromosomes (left; diploids: *G* = 23.7, *P* = 0.07; haploids: *G* = 12.0, *P* = 0.68) was correlated with the relative distance between the centromere and the chromosome midpoint in diploids (right), and this correlation differs significantly between ploidy levels (bootstrap *P* < 0.01). Centromere location was calculated as (*Lq* − *Lp*)/(*Lq* + *Lp*), where *Lq* is the length of the long arm and *Lp*is the length of the short arm. See Dataset S2 for a summary of variants by chromosome.

We found that substitutions but not indels occurred further from the centromere than expected by chance (Fig. 5B), but neither ploidy nor *RDH54* status affected the average distance of mutations from the centromere (all | *t* | < 0.99, all *P* > 0.32). Homozygous SNMs were especially likely to occur far from the centromere (Fig. 5B), which is expected if LOH is caused by mitotic crossing over, though non-crossover gene conversion has also been found to be more frequent further from the centromere (17). Homozygous variants occurred non-randomly among chromosomes (χ^2^ = 114, *P* < 10^−15^), and the distribution differs from that of heterozygous variants (χ^2^ = 143, *P* < 10^−15^). This pattern is driven almost entirely by an excess of homozygous variants on chromosome XII (21 of the 39 events, all in different lines; binomial test: *P* < 10^−12^), all distal to the rDNA tandem repeat region (Fig. S7), which is known to exhibit a high LOH rate (25, 26). Excluding events on chromosome XII, homozygous variants still occur further from the centromere than expected by chance (Wilcoxon and Kolmogorov-Smirnov tests: *P* < 0.01).

Previous studies have found evidence that mutation rates are correlated with the GC content of surrounding regions (17, 27). While we find no correlation between SNM rate and GC content (in accordance with ref. (5), we find that indels are associated with low GC content (in accordance with ref. (17), whereas MNMs are associated with high GC content (Fig. 5C). These associations diminish rapidly with distance from the focal site.

While the number of SNMs on a given chromosome was highly correlated with callable chromosome length (haploids: *r* = 0.96, *P* < 10^−8^; diploids: *r* = 0.96, *P* < 10^−8^), there was some evidence for residual variation (Fig. 5D), and we found that diploid SNM rates were significantly lower on chromosomes where the centromere is relatively distal, whereas this correlation was absent, and possibly reversed, in haploids (Fig. 5D). We reasoned that if this unexpected result reflects genuine mutation rate variation among chromosomes the pattern might also be evident in the locations of polymorphisms. We found that polymorphism rates by chromosome in wild and domesticated yeast isolates (28) were more strongly correlated with our diploid SNM rates (*ρ* = 0.79, 95% CI: 0.45 to 0.95, *P* < 0.001; Fig. S8), than our haploid SNM rates (*ρ* = −0.05, 95% CI: −0.50 to 0.41, *P* = 0.87), as expected if polymorphism reflects mutation rates and given that *S. cerevisiae* is generally diploid. As with SNMs in diploids, polymorphism rates tend to be negatively correlated with centromere location (*ρ* = −0.47,*P* = 0.066).

Examining the spatial pattern of SNMs within chromosomes, we found that the extent of spatial correlation between ploidy levels was itself negatively correlated with centromere location (*r* = −0.70, *P* < 0.01; Fig. S9), which is unlikely to occur by chance (5000 simulated datasets, *P* < 0.01). This suggests that the mutation rate in diploids is reduced in particular regions of chromosomes with distal centromeres. Additional possible explanatory factors are addressed in *SI Text*. The relative position of centromeres will affect the physical location of chromosome arms within the nucleus, as well as interactions between homologous loci (29−31), which might differentially affect haploid and diploid genome stability.

### Growth in Relation to Mutations

In our study, MA led to reduced growth in diploids but not haploids (Fig. 1C). This difference was driven, at least in part, by a particular class of mutations—aneuploidy events—that arose in diploid but not haploid lines (Fig. 3B). Accounting for these large-effect variants, including the presence of chromosome XI trisomy in the ancestral diploid *RDH54* + control, we modeled the effects of genome size and number of nonsynonymous nuclear and mt point mutations using mixed models (Fig. S10; see *SI Text* for details). We confirmed a highly significant effect of genome size change (χ^2^ = 291.37, *P* < 10^−15^) and detected an interaction between ploidy and point mutations (χ^2^ = 5.91, *P* < 0.05). Separate models for haploids and diploids showed a negative but nonsignificant effect of point mutations in haploids (χ^2^= 0.32, *P* = 0.57) and a significant negative effect in diploids (χ^2^ = 19.56, *P* < 10^−5^).

These results, as well as other modeling approaches (*SI Text*), indicate that point mutations had stronger negative effects on diploids than haploids, controlling for the effects of aneuploidy. It is possible that the spectrum of point mutations that accumulated in diploids had more severe fitness effects on average. Alternatively, given that we detect significant genetic variance in haploid MA lines, more mutations may have had beneficial effects in haploid lines (see (32–34). Finally, the greater fitness decline in diploids could arise if beneficial mutations are common at both ploidy levels but are more recessive than deleterious mutations on average. More work is needed to relate the molecular changes caused by spontaneous mutation to effects on fitness (e.g., (35).

## Discussion

Despite intense study of DNA replication and repair in yeast, the effect that ploidy has on the genome-wide rate and spectrum of mutations has not been investigated prior to this study. Populations of haploid and diploid yeast frequently show distinct evolutionary behaviour in experimental systems (15, 16, 36-38), but it is unclear what role ploidy-specific mutational variation plays in these differences. We find evidence that multiple dimensions of the mutation spectrum depend on ploidy state in an otherwise isogenic background. In particular, diploidy confers greater replication fidelity with respect to single-nucleotide changes and mitochondrial mutations, but diploids are also more prone to large-scale mutations and deleterious fitness effects.

While we accounted for rates of cell division and applied similar mutation-calling criteria in haploids and diploids, it is worth considering whether other differences between ploidy levels could have biased our mutation rate comparisons. To examine whether our coverage criteria might differentially affect haploid and diploid mutation rate estimates we re-estimated SNM and indel rates using progressively more-stringent cutoffs, and found no indication that mutation-calling criteria affected our results (Fig. S11).

Recessive lethal mutations can accumulate in diploids but not in haploids, which could lead us to underestimate the haploid mutation rate. Using the viability of spores from diploid MA lines, Zhu et al. (5) estimated the rate of recessive lethals as 3.2 × 10^−5^ per diploid genome per generation, involving SNMs and indels. Applying half this rate to haploids, we would have missed only ∼5 mutations among all of the haploid lines due to recessive lethality (a bias of <1%). Similarly, aneuploidy might not be observed in haploid lines if such changes are highly deleterious. However, aneuploidy for ten different chromosomes has been observed in haploid MA lines with mutator genotypes (7), and doubling times are similar to the wild type for at least some disomic haploids (39), suggesting that selection does not fully explain the lack of aneuploidy events in haploids.

Mutations might be expected to accumulate at a lower rate in haploids than diploids due to a greater effectiveness of selection against partially-recessive deleterious alleles. However, we found no evidence for selection on SNMs, and a consistent pattern of apparent selection across ploidy levels based on genic versus non-genic indel rates (Fig. S2). While the dearth of genic indels suggests purifying selection, mutations may not always arise at the same rate in genic and non-genic regions (40), complicating this interpretation.

We predicted that the rate of indels might be greater in haploids and *rdh54Δ* diploids owing to their reduced opportunity for homologous DNA repair, which did not prove to be the case (Fig. 2). We have lower statistical power with respect to indels, which are rare relative to SNMs. However, given the mean experiment-wide indel rate observed, we would detect a two-fold higher indel rate in haploids approximately 89% of the time at the α = 0.05 level. Similarly, given the mean diploid lines indel rate, we would detect a two-fold higher indel rate in *rdh54Δ* versus *rdh54* + lines approximately 85% of the time.

The lack of a difference in indel accumulation between ploidy levels or any interaction with *RDH54*status suggests that most small indels are the result of processes other than DSB repair, such as replication slippage (41), or that most DSBs are repaired when sister chromatids are available (3, 42). However, our results point to another possible effect of ploidy on DNA repair; the finding that replication timing affects mutation rates more strongly in haploid than diploid genomes suggests that the activity of error-prone DNA repair processes in late-replicating regions (specifically translesion synthesis, (23) is more important in haploids, which may explain why an effect of replication timing is detected in some studies (ref. (23) in haploids) but not others (ref. (43), in diploids). Replication timing may also explain the difference we observed in the spectrum of SNMs between ploidy levels, as this difference increased with replication timing (Fig. S12), with late replicating regions more prone to A-to-T and C-to-G transversions in haploids. Rather than increasing the rate of indels, the most obvious effect of *rdh54Δ* was to increase the rate of aneuploidy, suggesting that this gene plays a role in mitotic segregation.

LOH in diploids will convert some heterozygous mutant sites into sites homozygous for the ancestral allele, causing us to underestimate diploid point mutation rates. Using maximum likelihood to jointly estimate the rate of LOH and the true mutation rate (*SI Text*), we find that the rate of LOH [95% CI] is 7.92 [5.69, 10.65] × 10^−5^ per base pair per generation and that the observed diploid mutation rates are not substantially downwardly biased by LOH (3% bias; corrected diploid rates: SNMs: 2.97 [2.81, 3.15] × 10^−10^, indels: 2.09 [1.67, 2.57]×10^−11^]. These relative rates of mutation and LOH predict that few heterozygous sites will exist in equilibrium asexual populations (*SI Text*).

In our study we chose to examine diploids with the standard MATa/MATα genotype. Thus, results that we ascribe to ploidy differences may also stem from differences in MAT genotype. Although we did not detect an effect of MATa vs. MATα in haploids, it would be valuable to examine the mutational profiles of MATa/MATa and MATα/MATα diploids to clarify the role of MAT heterozygosity versus ploidy, *per se* especially given that MAT is known to affect repair pathway regulation (3) and competitive fitness (44).

We observed a much greater rate of mutation accumulation in mitochondrial sequences within haploid cells than diploid cells. In our study, nuclear relative to mt coverage does not depend on ploidy level (*t* = 1.62, *df* = 218, *P* = 0.11), suggesting that diploids have about twice the mt DNA of haploids (see qualitatively similar results in (45). Assuming sequencing and mapping are equally efficient for nuclear and mt DNA, we estimate that there are 3−4 sequenced mtDNA genomes per haploid cell, fewer than most previous estimates, which also indicate variation among strains and conditions (e.g., (45–47). However, we also see substantial variation in coverage across the mt genome, so coverage metrics may poorly estimate the number of mt genomes per cell. While we cannot exclude the possibility that a higher effective mt population size in diploids permitted more effective selection against new mutations, most of the mt mutations we observed were non-genic and presumably had minimal fitness effects. Instead, the factors causing a higher nuclear mutation rate in haploids may have an even greater impact on mt sequences, or haploids may incur a higher level of mt DNA damage. A study using a reporter gene approach found that mt microsatellites were 100 times more stable in diploids than in haploids (48), attributable to nuclear ploidy *per se* rather than *MAT*-specific gene expression or mt copy number (49). The effect of ploidy on mt genome maintenance clearly deserves further study.

In an experimental evolution context haploid populations of *S.cerevisiae* are sometimes found to attain a diploid state (50–52), but it is unclear to what extent this is due to a high rate of diploidization or a large selective advantage of diploidy. We observed no changes in ploidy, indicating that spontaneous diploidization does not occur at a high rate in the strain and experimental conditions we used. Assuming a rate of diploidization equal to our upper 95% confidence limit, a haploid population would incur >200 non-synonymous point mutations for every ploidy change event. However, our data also suggest that shifts to diploidy will increase the rate of large structural mutations, consistent with analyses of adaptive genome evolution in haploids and diploids (15, 16, 52). Ploidy shifts, which often seem to lack immediate benefits in yeast (50, 53), may therefore be favoured in some environments due to their effects on the mutation spectrum.

We find that the mutation rate and spectrum differs dramatically between the haploid and diploid forms of a common genetic background. Our results contribute to the growing understanding that the mutation process can vary in response to genetic and genomic context, influencing the distribution of new genetic variation among individuals and populations.

## Materials and Methods in Brief

For details on growth rate analyses, phenotyping, bioinformatics, and flow cytometry see *SI Materials and Methods*.

Isogenic haploid and diploid strains were generated starting with a single haploid cell, using single-colony bottlenecks throughout the procedure. The strain SEY6211 (*MAT* a, *ho*, *leu2-3 112*, *ura3-52*, *his3-Δ200*, *trp1-* Δ901, *ade2-101*, *suc2-Δ9*) was obtained from ATCC (Manassas, VA) and induced to switch mating type using a standard plasmid transformation protocol. *RDH54* in cells of each mating type via transformation, and confirmed by Sanger sequencing. *MAT* a and *MAT*á haploids was deleted (positions 383065 to 386180 on chromosome II), replaced with *KanMX* deletion. These strains were frozen as ancestral controls and plated to single colonies to begin the bottlenecking procedure.

MA was conducted on solid YPAD media on 6 cm diameter plates by bottlenecking each line to a single colony every 24 h, picking the nearest isolated colony to the center of each plate under 6.3 × magnification. Plates were incubated at C, and plates from the previous day stored at 4 were mated to generate diploids with and without the *RDH54* C as backup. Mitochondrial petite mutations result in very small 24 h colonies, and we used the backup plate if all colonies on a plate were petite, if individual colonies could not be distinguished, or if colonies were absent. Backup plates were required in 0.4% of transfers, with no significant differences among treatments.

Early in the experiment 13 putatively haploid *MAT*á *RDH54+* lines were found to be diploid due to mating immediately following the mating type switch during strain construction, leading to a mixed colony of *MAT*á and diploid cells; we lines by subdividing five existing lines confirmed to be haploid. We later found that the diploid *RDH54+* lines generated by controlled crossing, but not those obtained by continued to propagate these diploids and added 15 *MAT*á *RDH54+* unintentional mating, were trisomic for chromosome XI; furthermore, the ancestral diploid *RDH54+* strain (but not *rdh54Δ*) used in phenotype assays carries this trisomy (confirmed by Illumina sequencing).

Concurrent with bottleneck 100, each ancestral control genotype was thawed and plated in the same fashion as the MA lines. The next day a single colony of each MA line and three replicate colonies from each control plate were transferred to 3 mL of liquid media, grown for three days at 30°C, and frozen in 15% glycerol. These frozen cultures were subsequently used for DNA extraction and sequencing, flow cytometry, and phenotyping.

DNA was extracted from 10 mL saturated cultures using a standard phenol/chloroform method and quantified with fluorometry. 0.4 ng of DNA was used to construct libraries using the Illumina Nextera XT kit and protocol. Libraries were pooled such that diploid samples had twice the concentration of haploid samples (to give equivalent coverage per chromosome) and sequenced in a single Illumina NextSeq lane with paired-end 150-bp reads (average coverage per line before screening: haploid 24.5 ×, diploid 47.6 ×). Reads were aligned with BWA *mem* (54), and mutations in non-repetitive regions called using GATK HC (55), following recommended practices. We screened sites based on multiple metrics, and calculated mutation rates accounting for the number of callable sites in each line.

## Acknowledgements

We are grateful to A. Kuzmin for assistance with sequencing and transformations, A. Gerstein, M. Dunham and M. Whitlock for helpful discussion, A. Johnson for flow cytometry support, and C. Landry for providing plasmids. This work was supported by a Banting Postdoctoral Fellowship to NPS and NSERC Discovery grant RGPIN-2016-03711 to SPO.

